# Spatially Resolved Epithelial States Delineate Immunosuppressive Niches of Impaired Myeloid Antigen Presentation in PDAC

**DOI:** 10.64898/2026.07.28.741300

**Authors:** Cameron Walker, Ruohan Wang, Hadeesha Piyadasa, Brooks Benard, Carl Pelz, Jenny Eng, Kevin Hawthorne, Rosalie Sears, Andrew J. Gentles, Tyler Risom, Michael Angelo

**Author notes:** Correspondence should be addressed to MA or TR. These authors contributed equally to this work as corresponding authors (Tyler Risom, Michael Angelo).

## Abstract

Bulk transcriptomic classifiers stratify pancreatic ductal adenocarcinoma (PDAC) into classical and basal-like subtypes with prognostic and therapeutic relevance, yet increasing evidence indicates that these epithelial programs frequently coexist within individual tumors. How these intermediate states affect the local tumor microenvironment remains poorly defined. Here, we integrate multiplexed ion beam imaging (MIBI) with bulk RNA sequencing to resolve epithelial subtype identity at the level of spatially contiguous cancer nests and quantify their associated microenvironments. Across 47 primary tumor samples from 34 patients, we identified classical, intermediate, and basal cancer cell states at single-cell resolution and delineated discrete cancer nests with mixed or dominant subtype compositions. Distance-resolved spatial analysis reveals that basal-rich cancer nests are surrounded by locally immunosuppressive microenvironments characterized by reduced expression of MHC class II and co-stimulatory molecules in proximal myeloid cells, independent of myeloid abundance. These regions are enriched in fibroblast-dominated neighborhoods and distinct cell-cell interaction architectures. Using EcoTyper analysis of two independent bulk RNA-seq cohorts, including an OHSU discovery cohort (N = 277 patients) and TCGA as a validation cohort (N = 147 patients), we identified poor-prognosis tumor ecotypes enriched for basal epithelial states that similarly exhibited depleted myeloid antigen presentation signatures, linking spatial niche phenotypes to transcriptional ecotypes and patient outcomes. Together, these findings demonstrate that epithelial subtype programs in PDAC are organized at the level of spatially defined cancer nests and that basal cancer programs reside within localized niches of myeloid antigen presentation dysfunction, linking intratumoral architecture to immune suppression and clinical prognosis.

## Introduction

Pancreatic ductal adenocarcinoma (PDAC) is a highly lethal malignancy with limited therapeutic options (1). Transcriptomic profiling has been used in multiple classification frameworks to stratify PDAC into classical and basal-like molecular subtypes. Basal-like tumors are associated with worse survival, increased metastatic burden, chemoresistance, and poor response to FOLFIRINOX, whereas classical tumors exhibit relatively better outcomes (2–5). These observations have led to the clinical adoption of bulk RNA-based subtype classifiers, particularly Purity Independent Subtyping of Tumors (PurIST), to guide first-line treatment selection.

Although bulk subtyping has proven clinically useful, it assumes that tumors converge on a single dominant epithelial program. A growing body of single-cell and spatial studies now shows that this assumption is often incorrect (6–9). Cancer cells with classical and basal programs commonly coexist within the same PDAC tumor and even within the same cancer nest, defined here as a contiguous group of malignant epithelial cells (7,9,10). Single-cell RNA sequencing has demonstrated the presence of both programs in primary tumors that were classified as classical by bulk profiling, leading to the designation of a new “intermediate co-expressor” (IC) tumor cell state (7,9,11,12). These findings have been corroborated at the protein level by multiplexed immunofluorescence (mIF) studies which detected a KRT17+GATA6+ intermediate phenotype in over 90% of tumors (9).

Intriguingly, coexistence of classical and basal programs appears to be spatially patterned (7,10). Spatial transcriptomic and histologic analyses indicate that different regions of the same tumor can exhibit distinct epithelial subtype programs, raising the possibility that epithelial identity is regulated locally through a mixture of cytokine signaling, chemotactic extracellular signals, and metabolic remodeling at the epithelial-tumor interface (10,11,13,14). Recent work has begun to move this from correlation toward causation: coculture of patient-derived organoids with cancer-associated fibroblasts (CAFs) shifts tumor cells from classical toward basal transcriptional programs, implicating the fibroblast secretome as an active determinant of epithelial identity (15). However, spatial studies to date have characterized these relationships at the level of individual cells or of pathologist-selected regions, leaving unresolved how discrete cancer nests of different epithelial identity interact with their surrounding microenvironment.

Here, we address this knowledge gap by integrating high-resolution, high-dimensional spatial proteomics with bulk transcriptomic analysis to define how epithelial subtype identity is coupled to the local tumor microenvironment in PDAC. Using a 40-marker spatial proteomics panel that simultaneously captures epithelial lineage, immune, and stromal markers, we resolve classical, intermediate, and basal cancer cell states at single-cell resolution within intact tumor sections and delineate spatially contiguous cancer nests of distinct subtype composition. To characterize how nest composition relates to the local environment, we deploy a set of spatial analyses that quantify immune, stromal, and extracellular matrix features across defined-distance radial bands extending from the borders of cancer nests. This approach reveals that myeloid cells near basal-rich cancer nests express lower levels of proteins associated with antigen presentation, including reduced expression of major histocompatibility complex class II and associated co-stimulatory molecules. We extend these observations to independent, larger PDAC cohorts and show that bulk RNA-defined poor-prognosis ecotypes enriched for basal epithelial signatures similarly display depleted antigen presentation programs in the myeloid compartment. Together, these findings link tumor microenvironmental states to transcriptional ecotypes and patient outcome, demonstrating that basal tumor programs reside within niches characterized by suppressed myeloid antigen presentation.

## Results

### Multiplexed imaging resolves heterogeneous cancer cell subtypes in PDAC

To characterize the spatial organization of epithelial subtype programs and their associated microenvironments at single-cell resolution, we assembled a cohort of 47 tumor samples from 34 PDAC patients and performed multiplexed ion beam imaging by time-of-flight (MIBI-TOF) using a 40-marker antibody panel targeting epithelial lineage markers, immune cell markers, stromal markers, and functional proteins (**Fig. 1A-B**). In parallel, we leveraged two independent bulk RNA-seq cohorts: an Oregon Health & Science University (OHSU) discovery cohort (N = 277 patients) and t TCGA PAAD validation cohort (N = 147 patients) for transcriptomic ecotype analysis (**Fig. 1B**) (16,17).

**Fig. 1.**
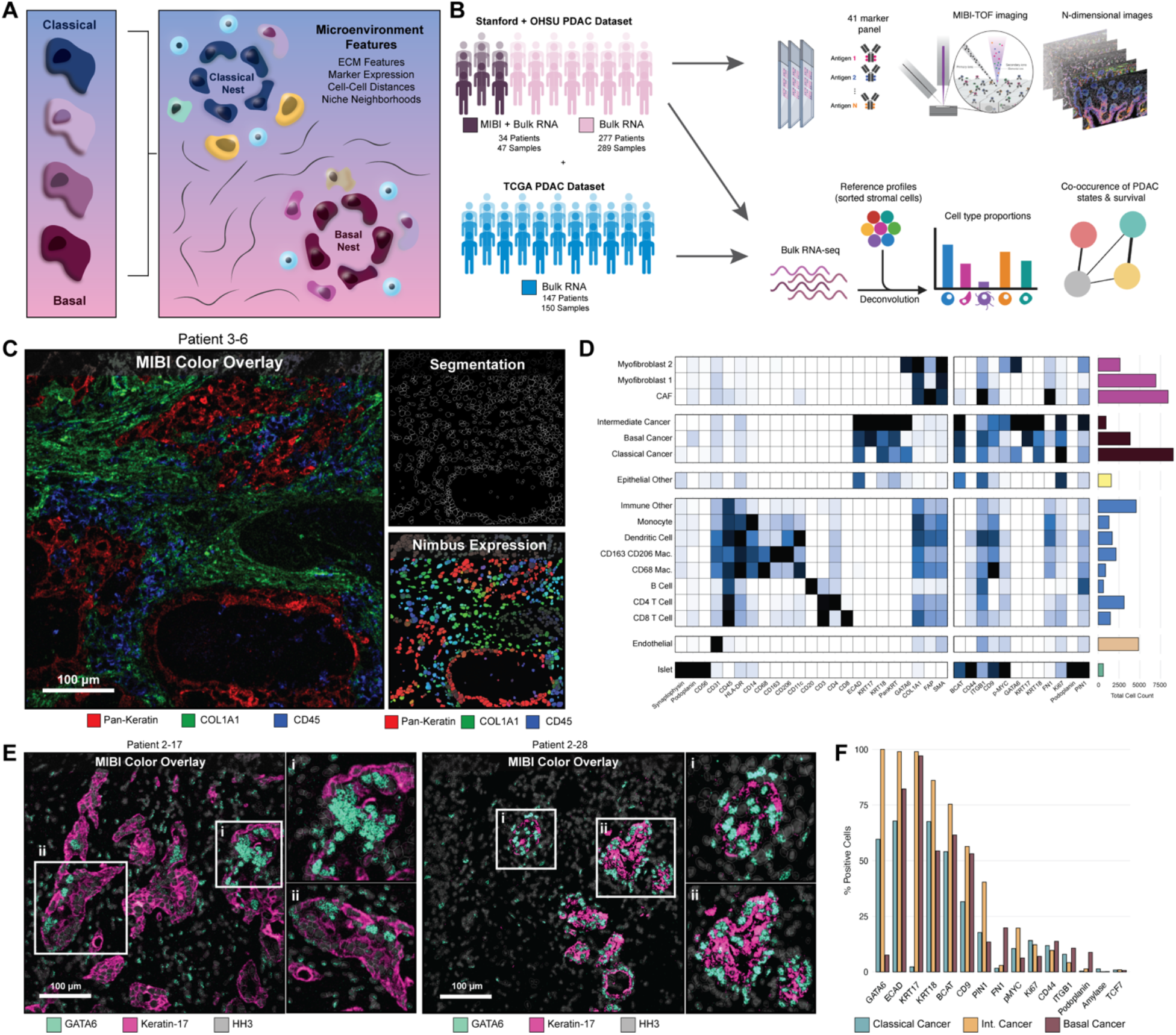
Multiplexed imaging reveals heterogeneous cancer nest subtypes in PDAC. (A) Overview of spatial analysis approach linking basal and classical cancer nest microenvironments. (B) Cohort design integrating MIBI-TOF (34 patients, 47 primary tumor cores) with bulk RNA-seq datasets (OHSU: 277 patients; TCGA: 147 patients). (C) Representative MIBI-TOF image with single-cell segmentation and NIMBUS marker prediction visualization. (D) Marker expression heatmap across NIMBUS-assigned cell types showing lineage-specific signatures. (E) Representative images demonstrating co-expression of basal-associated KRT17 and classical-associated GATA6 within a single nest. (F) Percent of classical, intermediate, and basal cancer subtypes positive for functional markers.

Following image acquisition and single-cell segmentation, we applied NIMBUS, a deep-learning-based marker prediction framework, to assign cell-level marker positivity across all 40 channels (**Fig. 1C**) (18). Cell type annotations were generated using FlowSOM clustering on NIMBUS-predicted marker intensities, yielding 17 distinct meta-clusters spanning cancer cells, fibroblasts, immune populations, and endothelial cells (19). A marker expression heatmap confirmed lineage-specific marker signatures across assigned cell types (**Fig. 1D**), and NIMBUS prediction maps for individual markers (CD68, Pan-Keratin, FAP) demonstrated concordance between predicted marker positivity and fluorescence overlay images (**Supplementary Fig. S1A, S1F**).

Within the cancer cell compartment, we classified cells into three epithelial subtypes based on the expression of established lineage markers: classical cancer cells (GATA6-high, KRT17-low), basal cancer cells (KRT17-high, GATA6-low), and intermediate cancer cells co-expressing both markers. Representative MIBI-TOF images demonstrated that basal-associated KRT17 and classical-associated GATA6 frequently co-localized within individual cancer nests and, in some cases, within the same cell (**Fig. 1E**). Single-cell co-expression analysis confirmed a continuum of GATA6 and KRT17 expression across subtypes, with intermediate cells occupying the transitional zone (**Supplementary Fig. S1D-E**). We further quantified the functional marker expression across the three cancer subtypes. Basal cancer cells exhibited elevated positivity for adhesion (CD44, ITGB1) and immune-modulatory markers compared to classical cancer cells, whereas classical cancer cells showed higher E-cadherin (ECAD) and Ki67 expression (**Fig. 1F**). The composition of cancer cell subtype varied across patients and disease stages (**Supplementary Fig. S1B-C**), with most tumors harboring mixtures of all three subtypes.

### Spatially contiguous cancer nests exhibit distinct subtype composition and E-Cadherin expression

Having established single-cell cancer subtype annotations, we sought to define the spatial unit of epithelial identity by identifying spatially contiguous cancer nests within each tissue section. We first generated tumor compartment masks based on pan-keratin expression to distinguish tumor regions from the surrounding stroma (**Fig. 2A-C**). Within the tumor compartments, spatially contiguous cancer cells were grouped into discrete cancer nests by connected component analysis of the PanKRT-derived tumor compartment masks (see Methods) (**Fig. 2D**). Across the cohort, we identified 204 unique cancer nests and quantified the proportion of classical, intermediate, and basal cancer cells in each group. K-means clustering of the nest-level subtype proportions, with the number of clusters (k = 3) selected by silhouette analysis (maximal average silhouette width at k = 3; **Supplementary Fig. S2C**), resolved into three distinct groups: classical-rich nests (predominantly classical cancer cells), intermediate-rich nests (enriched for intermediate cells), and basal-rich nests (predominantly basal cancer cells) (**Fig. 2E**). This nest-level classification captured substantial within-tumor heterogeneity, as many individual tumors contained nests belonging to more than one cluster.

**Fig. 2.**
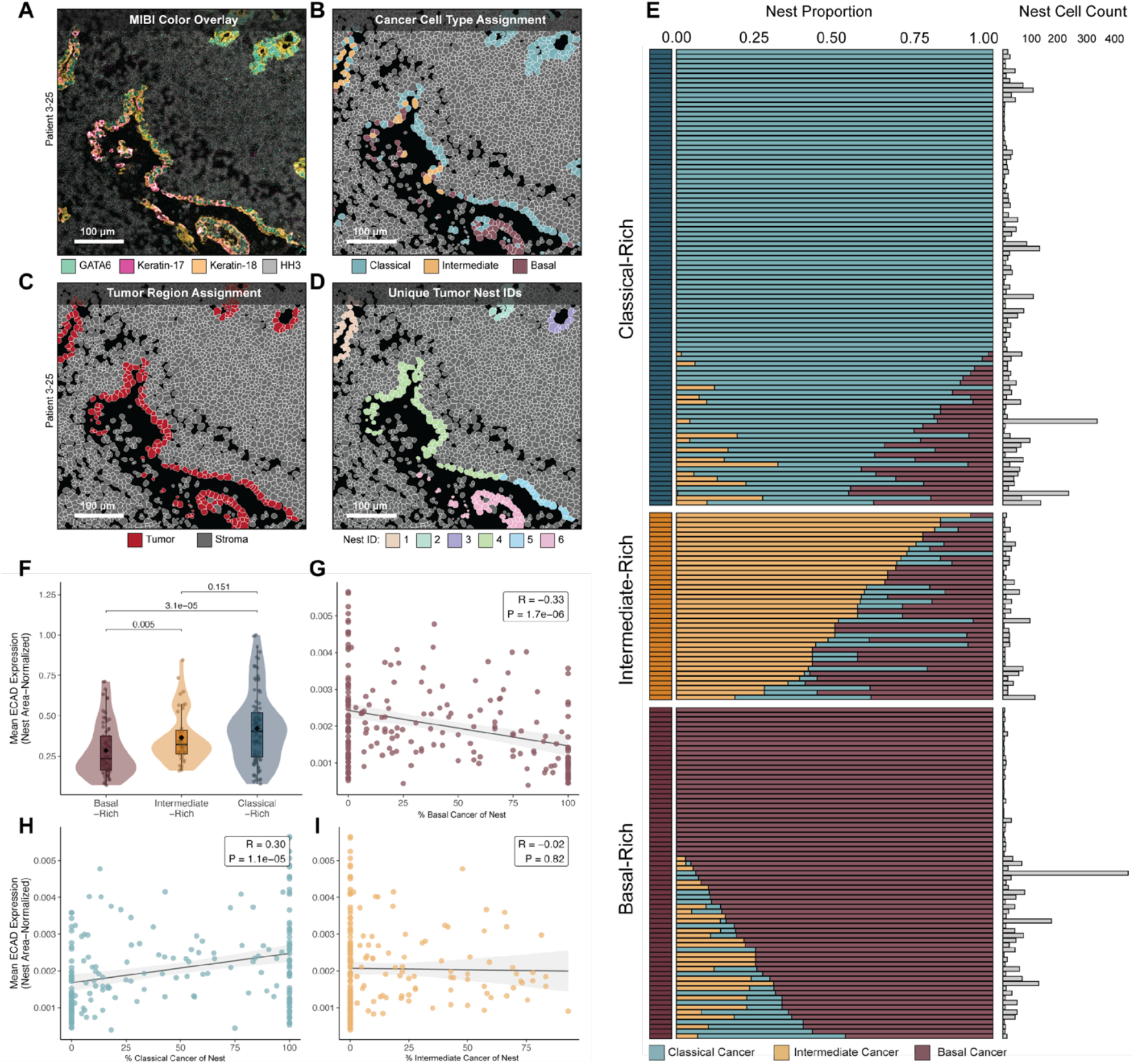
Tumor nest identity is linked to ECAD expression. (A) MIBI-TOF color overlay. (B) Cancer cell type assignment. (C) Tumor compartment masks distinguishing tumor from stroma. (D) Unique cancer nest identification within tumor regions. (E) Cancer nest composition across the cohort, clustered into three groups (classical-rich, intermediate-rich, basal-rich) by k-means clustering. (F) Mean ECAD expression across nest clusters. (G–H) Correlation between nest-level ECAD expression and cancer subtype composition.

Next, we investigated whether the cancer nest subtype composition was associated with adhesion molecule expression. The mean ECAD expression, normalized by nest area, was significantly higher in classical-rich nests compared to basal-rich nests (Wilcoxon rank-sum test with Bonferroni correction, P < 0.0001; **Fig. 2F**). At the nest level, ECAD expression was negatively correlated with the proportion of basal cancer cells (Pearson R = -0.33, P = 1.7 × 10^-6^; **Fig. 2G**) and positively correlated with the proportion of classical cancer cells (Pearson R = 0.30, P = 1.1 × 10⁻⁵; **Fig. 2H**). These findings are consistent with the known role of E-cadherin loss in epithelial-to-mesenchymal transition and basal-like programs in PDAC, and validate the biological coherence of our nest-level subtype classification (20).

### Distance-based spatial analysis reveals depleted antigen presentation near basal-rich nests

To systematically characterize the microenvironment surrounding cancer nests with different epithelial compositions, we developed a distance-based spatial feature-extraction framework. For each unique cancer nest, we computed the microenvironmental features within concentric distance bands extending outward from the nest border: 0-25 μm, 25-50 μm, 50-100 μm, 100-200 μm, 200-300 μm, 300-400 μm, and greater than 400 μm (**Fig. 3A**). Nearly 250 features, including cell type densities, cell type ratios, functional marker positivity across stromal and immune populations, and 46 extracellular matrix (ECM) features derived from COL1A1 fiber segmentation masks, were quantified within each concentric distance band surrounding each cancer nest. Representative images from three patients illustrate the spatial analysis workflow, showing distance band overlays along with COL1A1 segmentation, cell type ratio maps, and functional marker positivity maps (**Fig. 3B**).

**Fig. 3.**
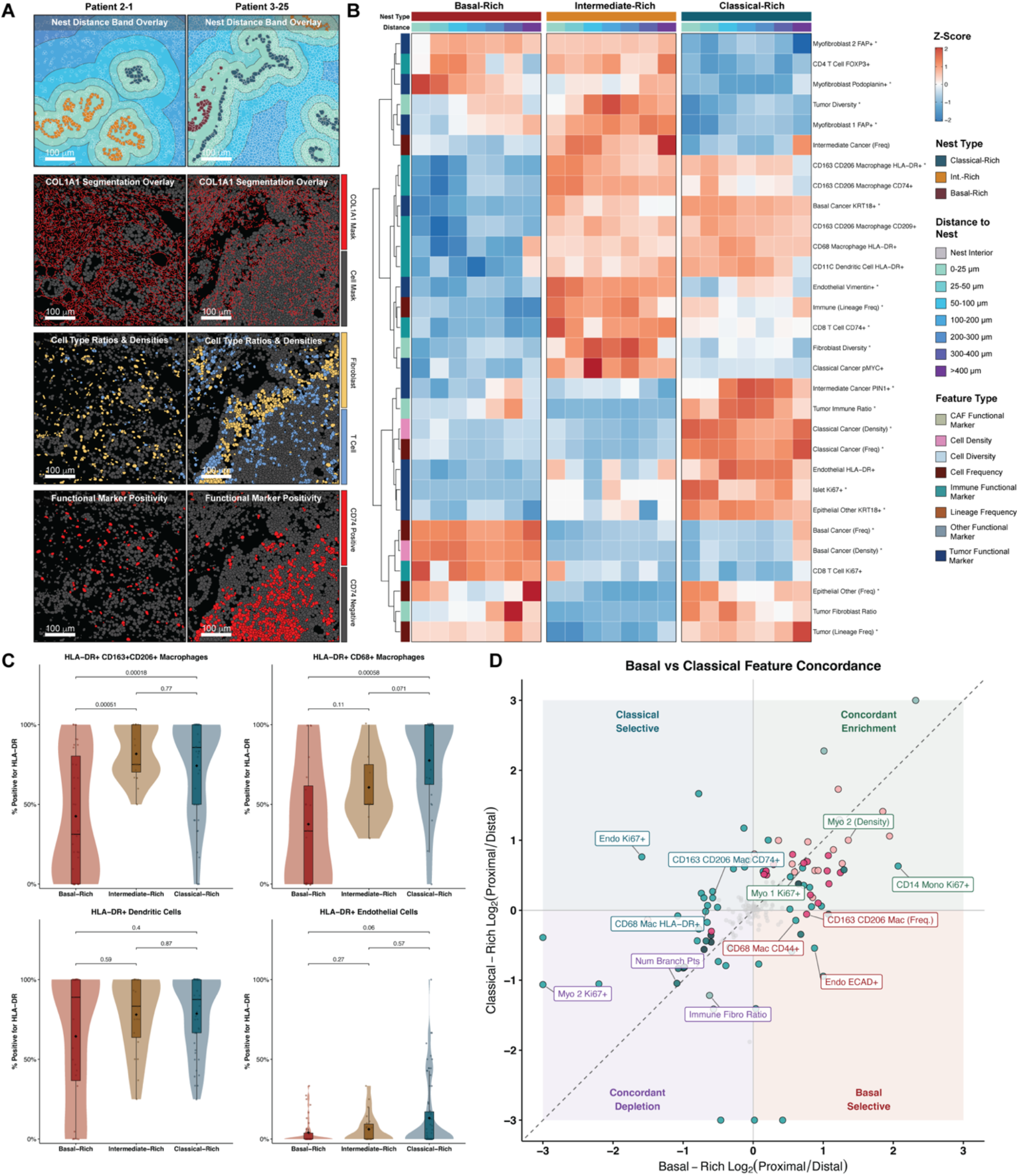
Distance-based quantification of spatial features reveals depleted HLA-DR and CD74 expression proximal to basal-rich nests. (A) Schematic of spatial feature extraction approach. Image features are computed at defined distance bands from each unique cancer nest border (0–25 μm, 25–50 μm, 50–100 μm, 100–200 μm, 300–400 μm, > 400 μm). Representative MIBI-TOF images from three patients illustrating the spatial analysis workflow. (B) Heatmap of significantly enriched spatial features by distance to each cancer nest type. Axis colored by feature type. (C) HLA-DR marker positivity in myeloid and endothelial cells within 0–50 μm of each cancer nest type. (D) Basal vs. Classical feature concordance plot comparing proximal enrichment profiles (dots colored by feature type).

We computed z-scored spatial features across all nests stratified by nest type and distance band and visualized significantly enriched or depleted features in a heatmap (**Fig. 3C**). This analysis revealed striking differences in the stromal microenvironment proximal to basal-rich versus classical-rich nests. Several broad patterns have emerged. Basal-rich nests were surrounded by regions enriched for CAFs and proliferating (Ki67+) CD8 T cells, particularly within the 0-50 µm distance band. Both patterns are consistent with a recent imaging mass cytometry study of PDAC, which identified myofibroblastic CAFs and T cells as the populations most frequently adjacent to basal tumor cells (15). Their recovery in an independent cohort on a different imaging platform supports the biological validity of our nest-level classification. Per-nest quantification confirmed that fibroblast density, myeloid cell density, and proportion of FOXP3+ regulatory T cells differed significantly across nest types in the proximal zone (**Supplementary Fig. S3A–C**). Comparison of proximal versus distal feature enrichment further identified CAF subsets as among the most strongly enriched populations near basal-rich nests (**Supplementary Fig. S3D–H**). In contrast, classical-rich nests were associated with distinct ECM organizational features and relatively preserved immune marker expression in the proximal stroma. Intermediate-rich nests exhibited a largely intermediate phenotype but also displayed unique features, including endothelial HLA-DR expression.

To directly assess the antigen presentation capacity of myeloid cells near the cancer nests of different subtypes, we quantified functional marker positivity in stromal cells within 0-50 μm of each nest type. HLA-DR positivity was significantly reduced in CD68 macrophages and CD14 monocytes proximal to basal-rich nests compared to classical-rich nests (**Fig. 3C**). CD74, the invariant chain required for MHC class II peptide loading, showed a concordant pattern of reduced expression near basal-rich nests (**Fig. 3E**). These differences were not explained by differential myeloid cell abundance, as macrophage and monocyte densities did not differ significantly across nest types in the proximal distance band (**Supplementary Fig. S3B**).

To further characterize features that distinguish the microenvironment immediately adjacent to cancer nests from the more distal stroma, we performed a proximal (< 50 μm from cancer nest) versus distal (> 200 μm from cancer nest) enrichment analysis for each nest type. Differential feature enrichment revealed that basal-rich nests were proximally enriched for CAF1 densities, tumor-stromal ratios, and proliferating fibroblast subsets, while being depleted for endothelial cells, certain immune populations, and ECM organizational features at the proximal zone while Intermediate-Rich and classical-rich nests showed partially overlapping but distinct proximal enrichment profiles (**Supplementary Fig. S3F-H)**. A concordance analysis comparing proximal enrichment between basal-rich and classical-rich nests identified features that were selectively enriched near one nest type but not the other (**Fig. 3D**). Features in the lower-right quadrant were basal-rich-selective (enriched only near basal-rich nests), whereas those in the upper-left quadrant were classical-rich-selective. Basal-rich-selective features included elevated tumor-fibroblast ratios and CAF marker positivity, whereas classical-rich-selective features included CD8 T cell densities and specific ECM metrics.

To complement these distance-based analyses, we applied QUICHE, an independent spatial method that defines local cellular niches based on neighbor composition and tests for differential enrichment proximal versus distal to the cancer nests (21). QUICHE confirmed that CAF-dominated neighborhoods composed predominantly of CAF1 and CAF2 fibroblasts were significantly enriched proximal to basal-rich nests, whereas classical-rich nests were surrounded by neighborhoods with greater immune cell representation, including CD4 T cells and endothelial cells (**Supplementary Fig. S4**). Intermediate-rich nests showed niches enriched for CD163/CD206 macrophages co-occurring with mixed cancer cell populations. In parallel, we characterized extracellular matrix organization by processing COL1A1 images through TWOMBLI to extract 46 ECM features per field of view (22) (**Supplementary Fig. S5A**). ECM features varied according to tumor grade, with higher-grade tumors exhibiting increased porosity and decreased fiber orientation coherence (**Supplementary Fig. S5B–E**), and myofibroblast-associated fibroblast subsets showed the strongest correlation with fiber organizational metrics (**Supplementary Fig. S5F**). Notably, CAF density was positively correlated with collagen fiber length, whereas myofibroblast-1 and myofibroblast-2 densities were more strongly associated with fiber fractality and topological organization metrics (**Supplementary Fig. S5I**; red indicates a statistically significant correlation). Unsupervised clustering of ECM profiles identified two distinct ECM states associated with tumor grade and myofibroblast enrichment, although ECM cluster membership did not independently predict overall survival or was associated with a specific epithelial lineage (**Supplementary Fig. S5G–K**).

### EcoTyper identifies prognostic tumor ecotypes across independent PDAC cohorts

To determine whether the spatial phenotypes observed at the cancer nest level corresponded to transcriptionally defined tumor states at the bulk level, we applied EcoTyper to two independent bulk RNA-seq cohorts (16,23). EcoTyper decomposes bulk RNA-seq data into cell-type-specific expression programs, identifies transcriptionally defined cell states within each cell type, and groups co-occurring cell state patterns into ecotypes across patients (**Fig. 4A**).

**Fig. 4.**
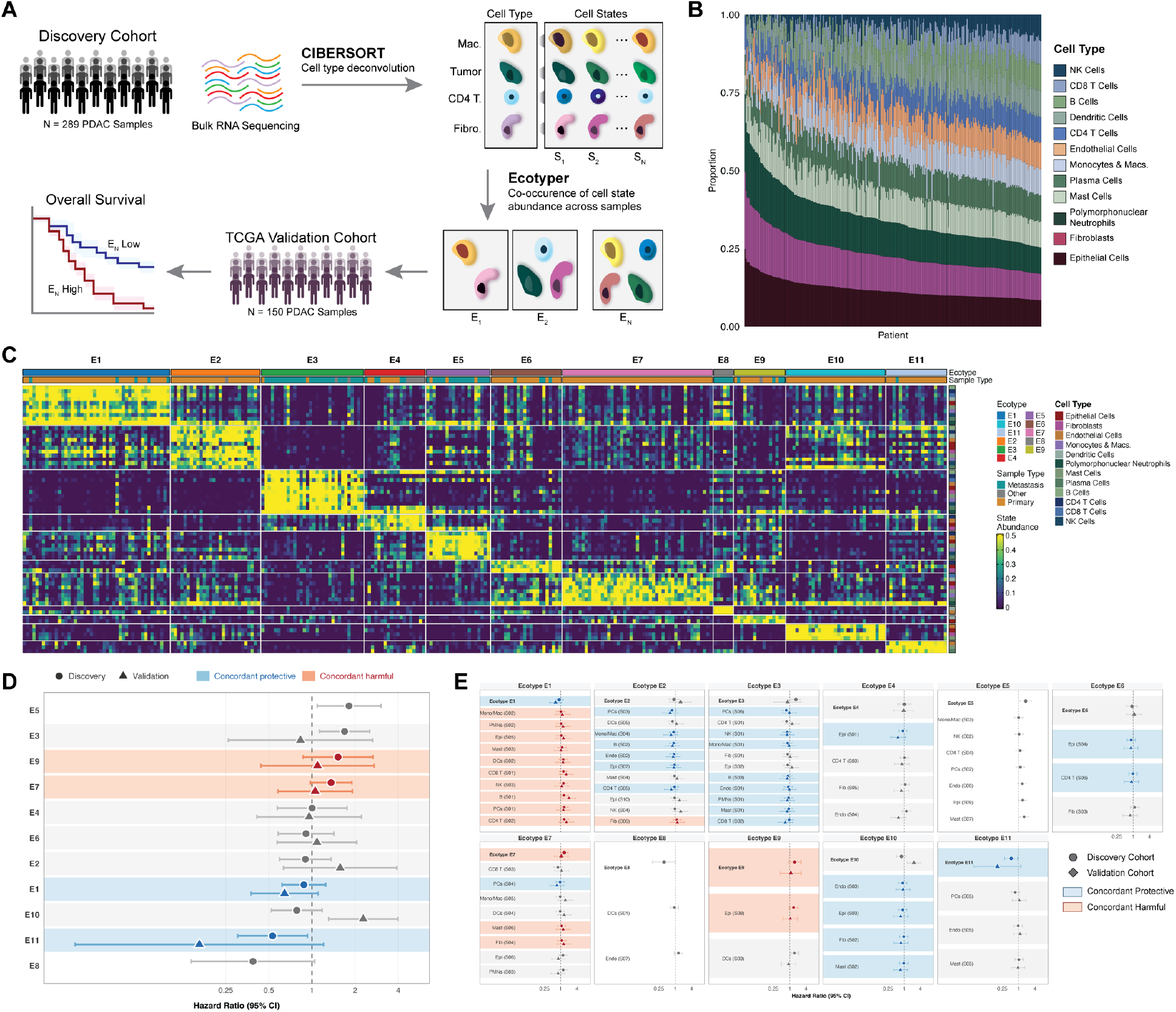
EcoTyper identifies prognostic tumor ecotypes validated across independent PDAC cohorts. (A) Schematic of EcoTyper workflow. Bulk RNA sequencing is decomposed into cell-type-specific expression programs and transcriptionally defined cell states, which are filtered by recovery in two independent PDAC scRNA-seq datasets and grouped into co-occurring ecotypes. (B) Cell type composition across discovery cohort patients estimated by CIBERSORTx deconvolution. (C) Heatmap of cell state abundances across discovery cohort samples. (D) Forest plot of ecotype-level hazard ratios for overall survival from assignment analysis. Blue shading indicates concordant protective association; orange shading indicates concordant harmful association. (E) Cell state-level hazard ratios within each ecotype.

In the discovery cohort (N = 277 PDAC patients) (16), the cell-type-specific expression profiles for 12 cell types were extracted and decomposed into transcriptionally defined cell states (**Fig. 4B**). EcoTyper decomposition identified multiple cell states per cell type and grouped patients into 11 ecotypes (E1–E11) based on co-occurring cell state patterns (**Fig. 4C**). To evaluate the prognostic association of each ecotype, we computed hazard ratios with 95% confidence intervals (CIs) comparing patients assigned to each ecotype versus all others (Cox proportional hazards regression), and identified prognostic ecotypes by directional concordance across the two independent cohorts (**Fig. 4D**). Four ecotypes showed concordant prognostic associations across both the discovery and TCGA validation cohorts: E1 and E11 were concordantly protective (HR < 1 in both cohorts), whereas E7 and E9 were concordantly harmful (HR > 1 in both cohorts). Cell state-level hazard ratios within each ecotype further delineated the specific cell states that drove the prognostic associations (**Fig. 4E**).

### Poor-prognosis ecotypes exhibit reduced MHC class II expression and basal-shifted epithelial states

We next asked whether the concordant poor-prognosis ecotypes (E7 and E9) (**Fig. 5A**) recapitulated the myeloid antigen presentation deficits observed spatially near basal-rich cancer nests. To do so, we compared the expression of MHC class II molecules and co-stimulatory genes between the good- and poor-prognosis ecotype groups. To account for potential differences in macrophage abundance, we normalized antigen presentation gene expression to the Ecotyper-estimated macrophage fraction (summed abundance of monocyte/macrophage states). Notably, none of the myeloid markers tested, CD68, CD163, or MRC1, were differentially expressed between the ecotype groups (all P > 0.2; **Supplementary Fig. S6**), indicating that the total macrophage abundance did not differ between the good- and poor-prognosis ecotypes.

**Fig. 5.**
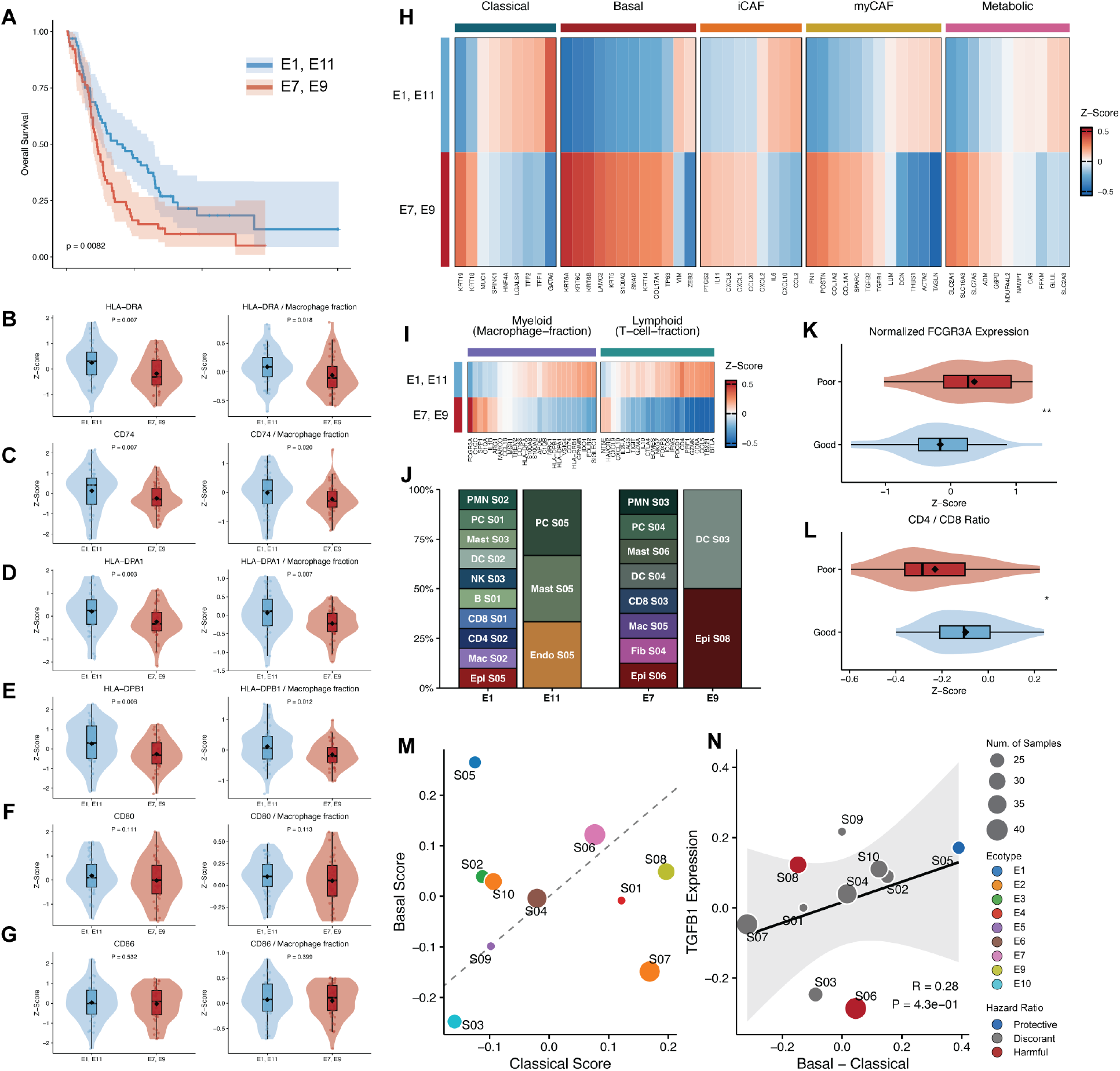
Poor-prognosis ecotypes exhibit reduced MHC class II expression and basal-shifted epithelial states in OHSU discovery cohort. (A) Kaplan-Meier survival analysis comparing concordant good-prognosis ecotypes (E1, E11) versus concordant poor-prognosis ecotypes (E7, E9). (B–G) MHC class II and co-stimulatory molecule expression in good- and poor-prognosis ecotypes normalized by macrophage fraction in ecotype: HLA-DRA (B), CD74 (C), HLA-DPA1 (D), HLA-DPB1 (E), CD80 (F), CD86 (G). (H) Gene expression signatures comparing good- and poor-prognosis ecotypes. (I) Immune cell marker expression normalized by lineage abundance. (J) Cell-state composition of each prognostic ecotype. Bars enumerate the transcriptionally defined cell states that co-occur to define the indicate ecotype. Segment height reflects that state’s share of ecotype defining cell states (K) Normalized FCGR3A expression. (L) CD4/CD8 T cell ratio. (M) Epithelial cell states positioned by basal and classical signature scores. (N) Correlation of basal-classical signature score and TGFB1 transcript abundance.

HLA-DRA expression was significantly reduced in poor-prognosis ecotypes, both as raw z-scored expression and after normalization to the EcoTyper macrophage fraction (**Fig. 5B**). CD74, the invariant chain required for MHC class II peptide loading, showed concordant reduction (**Fig. 5C**). Additional MHC class II genes, including HLA-DPA1 and HLA-DPB1, were similarly depleted (**Fig. 5D–E**). The co-stimulatory molecules CD80 and CD86, which facilitate T cell activation during antigen presentation, also showed reduced expression in poor-prognosis ecotypes (**Fig. 5F–G**), consistent with impaired antigen presentation capacity. The combination of unchanged macrophage marker expression with significantly reduced AP gene expression between ecotype groups indicates that the antigen presentation deficit reflects a per-cell functional difference in the myeloid compartment, rather than a difference in macrophage abundance or polarization composition.

Gene expression signature analysis confirmed that good-prognosis ecotypes (E1 and E11) were enriched for classical epithelial signatures, while poor-prognosis ecotypes (E7 and E9) exhibited predominantly basal-like signatures (**Fig. 5H**). Immune cell marker expression, normalized to EcoTyper-estimated lineage fractions (the macrophage fraction for myeloid markers and the T-cell fraction for lymphoid markers), further revealed that poor-prognosis ecotypes displayed reduced myeloid antigen presentation markers alongside altered lymphoid compartments (**Fig. 5I**). Cell state composition analysis revealed that the good- and poor-prognosis ecotypes were assembled from distinct sets of co-occurring cell states (**Fig. 5J**). Poor-prognosis ecotypes were anchored by basal-associated epithelial states (Epi S06 in E7, Epi S08 in E9) together with specific stromal and myeloid states (Fib S04, Mac S05), whereas good-prognosis ecotypes comprised classical-leaning epithelial states (Epi S05) along with a broader complement of lymphoid states (CD4, CD8, B, and NK cell states in E1) and a stromal-dominated architecture (PC S05, Mast S05, Endo S05 in E11). (**Fig. 5J).** Poor-prognosis ecotypes also exhibited increased FCGR3A (CD16) expression (P < 0.01; **Fig. 5K**) and altered the CD4/CD8 T cell ratios (P < 0.05; **Fig. 5L**), suggesting broader immune dysregulation beyond the myeloid compartment.

To characterize these immune and stromal differences at the pathway level, we computed gene signature scores for T-cell function, fibroblast polarization, and macrophage activation states across the ecotype groups (**Supplementary Fig. S7**). Poor-prognosis ecotypes exhibited elevated Tell exhaustion signatures with a reduced effector-to-exhaustion ratio (**Supplementary Fig. S7A–C**), consistent with a dysfunctional adaptive immune compartment. Fibroblast programs also differed, with poor-prognosis ecotypes showing a shifted iCAF-to-myCAF ratio (**Supplementary Fig. S7D–F**). In the myeloid compartment, poor-prognosis ecotypes displayed a reduced inflammatory-to-regulatory macrophage signature ratio (**Supplementary Fig. S7G–I**), concordant with the observed depletion of antigen presentation gene expression, indicating a coordinated skew toward immunoregulatory myeloid programs.

To directly link epithelial cell states to the basal-classical axis, we scored each EcoTyper-defined epithelial cell state for the basal and classical gene signatures. Epithelial states associated with poor prognosis (S05, S06) mapped to the basal-like quadrant of the signature space, whereas states associated with favorable outcomes occupied the classical quadrant (**Fig. 5M**). Notably, TGFB1 transcript abundance was not significantly correlated with the basal-classical signature score (Pearson R = 0.28, P = 0.45; Fig. 5N), suggesting that the association between basal programs and suppressed antigen presentation may not simply be mediated by TGF-β signaling.

Taken together, these bulk transcriptomic analyses demonstrate that the spatial relationship between basal epithelial programs and myeloid antigen presentation dysfunction observed by MIBI-TOF is recapitulated at the bulk transcriptional level across independent, larger PDAC cohorts. Poor-prognosis ecotypes are characterized by the co-occurrence of basal-shifted epithelial states with depleted MHC class II expression in the myeloid compartment, linking the tumor microenvironmental architecture to transcriptional ecotypes and clinical outcomes.

## Discussion

In this study, we investigated the local interaction between tumor nests and stroma using patient-matched spatial proteomics and bulk RNA sequencing to determine how epithelial subtypes and microenvironmental states are spatially related. Basal differentiation has been linked to poor patient outcomes in a large corpus of research using bulk analytical approaches (2,3). The spatially resolved approach used here allowed us to expand on these findings by extending the epithelial subtype assignment to individual tumor cells. We found that the epithelial subtype varies substantially in most PDAC tumors and that this variation is spatially organized at the level of individual cancer nests. Furthermore, subtype composition of cancer nests is linked to focal differences in myeloid function in the local TME. These findings bridge the bulk transcriptomic classification of PDAC subtypes and the spatial heterogeneity that exists within individual tumors.

The identification of intermediate co-expression cancer cells in PDAC that express both classical and basal gene programs has progressed through several stages. Single-cell transcriptomic and multiplexed immunofluorescence studies first identified cells co-expressing basal and classical markers in PDAC tumors and established that tumor cell identity exists on a phenotypic continuum between classical and basal differentiation. Prior protein-level characterizations of these intermediate states have defined this population using a small number of lineage markers, establishing the co-expression of GATA6 with KRT17 as definitive for the IC state. Here, with the breadth of the spatial proteomics panel, we expanded this signature, showing that the IC phenotype co-expresses GATA6, KRT17, CD9, PIN1, and Phospho-c-MYC and frequently co-exists within single tumor nests with both classical and basal cancer cells.

Our approach of delineating spatially contiguous cancer nests and quantifying microenvironmental features at defined distance bands provides a framework for studying how epithelial identity interfaces with local stroma *in situ*. Basal-rich cancer nests are surrounded by stroma enriched for cancer-associated fibroblasts and myeloid cells lacking HLA-DR and CD74, a phenotype consistent with impaired MHC class II-mediated antigen presentation. Critically, this depletion was not driven by reduced myeloid cell abundance near basal nests, indicating that the antigen presentation deficit reflects a qualitative difference in the myeloid functional state rather than the simple exclusion of immune cells from the tumor-stroma interface. QUICHE spatial niche analysis independently confirmed that the immediate neighborhoods of basal-rich nests ere dominated by fibroblast-rich niches, in contrast to the more immunologically diverse niches surrounding classical-rich nests (**Supplementary Fig. S3**).

These observations align with the emerging literature linking basal-like PDAC programs to immunosuppressive tumor microenvironments. Basal-like tumors are associated with increased stromal activation, myofibroblast polarization, and resistance to immunotherapy (24). Our findings demonstrate that these immunosuppressive phenotypes are locally concentrated around nests with basal epithelial identity, rather than being distributed uniformly across the tumor. Functional evidence that CAFs induce a classical-to-basal shift in patient-derived organoids (15) favors a model in which the fibroblast-rich niche actively shapes epithelial identity, and proximal CAF enrichment around basal-rich nests supplies the corresponding architecture in human tissue. What remains unresolved is whether the myeloid antigen presentation deficit is downstream of the same fibroblast program or reflects an independent axis originating from the basal cancer cells themselves; distinguishing these possibilities will require longitudinal and perturbation-based studies.

The extension of our spatial observations to bulk transcriptomic data through EcoTyper analysis strengthens the generalizability of these findings. Across two independent cohorts totaling 424 PDAC patients, we identified poor-prognosis ecotypes enriched for basal epithelial cell states that concurrently exhibited depleted MHC class II and co-stimulatory gene expression in the myeloid compartment. The convergence between spatially defined microenvironmental phenotypes (observed at the protein level by MIBI) and transcriptionally defined ecotypes (derived from bulk RNA-seq) suggests that the relationship between basal programs and myeloid state is a consistent feature of PDAC biology rather than an artifact of any single measurement platform.

In summary, we demonstrated that PDAC epithelial subtype programs are organized at the cancer nest level and that basal-enriched nests reside within spatially defined niches of myeloid antigen presentation dysfunction. By linking these spatial phenotypes to transcriptional ecotypes and patient outcome across independent cohorts, our findings provide a framework for understanding how intratumoral epithelial heterogeneity is organized with respect to the local immune landscape in PDAC and identify myeloid antigen presentation as a candidate therapeutic axis in basal-enriched disease because it is concentrated in spatially discrete niches rather than distributed uniformly, may be missed by bulk subtype–guided treatment selection and warrants spatially informed therapeutic strategies.

This study has several limitations. First, our MIBI cohort of 34 patients was modest in size. Larger imaging cohorts are needed to fully capture the heterogeneity of cancer nest-stroma interactions in PDAC. Second, the 40-marker spatial proteomics panel, while comprehensive for major lineage and functional markers, does not capture the full breadth of the antigen presentation machinery or fibroblast states, including myCAFs and iCAFs, which may mediate the observed phenotypes. Single-cell RNA sequencing of micro-dissected basal-rich and classical-rich regions would provide complementary molecular resolution. Third, our EcoTyper analysis operates on cell-type-specific gene subsets of bulk RNA-seq data, which provides population-level estimates of cell states, but cannot resolve spatial relationships. The concordance between the MIBI-TOF and EcoTyper findings is encouraging, but does not establish a direct spatial correspondence between the two platforms.

## Methods

### Patient cohort and tissue acquisition

A total of 47 primary PDAC tumor cores (**Supplementary Table S1**) were collected from 34 patients who underwent surgical resection at OHSU. Clinical data, including tumor stage, tumor grade, treatment history, and survival outcomes were recorded. Patient tissues and data were acquired with informed consent aligned with the Declaration of Helsinki and were obtained from the Oregon Pancreas Tissue Registry under theOregon Health & Science University IRB protocol #3609.

### MIBI-TOF imaging

Formalin-fixed, paraffin-embedded (FFPE) tissue sections were stained with a panel of 40 metal-conjugated antibodies (**Supplementary Table S2**) targeting epithelial markers (Pan-Keratin, GATA6, KRT17, KRT18, E-Cadherin, Synaptophysin, phospho-MYC, BCAT, Amylase, PIN1), immune markers (CD3, CD4, CD8, CD45, CD20, CD14, CD68, CD74, CD163, CD206, CD209, CD56, CD11c, HLA-DR, FOXP3, Calprotectin, Ki67), stromal markers (FAP, SMA, Podoplanin, COL1A1, Vimentin, Fibronectin), functional and adhesion markers (CD9, CD31, CD44, ITGB1, ECAD), and additional markers (Chymotrypsin, TCF7, CHP). Tissue sections were imaged on the MIBI-TOF instrument at a resolution of approximately 500 nm per pixel, generating multiplexed ion images across all 40 channels for each field of view (FOV).

### Image processing and single-cell segmentation

Raw MIBI-TOF images were preprocessed using standard compensation and noise-removal pipelines. Single-cell segmentation was performed using Mesmer/DeepCell to generate cell masks for each FOV (25). Cell-level marker intensities were extracted by averaging the pixel-level signals within each cell mask across all 40 channels.

### NIMBUS marker prediction and cell type annotation

To assign cell-level marker positivity, we applied NIMBUS, a deep-learning-based framework that predicts the probability of marker positivity for each cell based on the local image context (18). NIMBUS predictions were used as inputs for the cell-type annotations. FlowSOM clustering was performed on NIMBUS-predicted marker intensities using a 4 × 6 self-organizing map grid (24 initial clusters), which were subsequently merged into 17 cell meta-clusters based on marker expression profiles: Classical Cancer, Basal Cancer, Intermediate (co-expressor) Cancer, Epithelial Other, CAF1, CAF2, CAF3, CD8 T Cell, CD4 T Cell, B Cell, CD11c Dendritic Cell, CD68 Macrophage, CD163/CD206 Macrophage, CD14 Monocyte, Immune Other, Endothelial, and Islet cells(17).

### Cancer cell subtype classification

Cancer cells were classified into three subtypes based on the NIMBUS-predicted expression of GATA6 and KRT17: classical cancer (GATA6-high, KRT17-low), basal cancer (KRT17-high, GATA6-low), and intermediate cancer (co-expression of both markers). Subtype assignments were validated by examining single-cell co-expression distributions and log2-transformed GATA6/KRT17 expression ratios.

### Tumor compartment segmentation and cancer nest identification

Tumor compartment masks were generated based on pan-keratin (PanKRT) signal intensity combined with cell segmentation overlays. Briefly, PanKRT images were Gaussian-smoothed (σ = 10 pixels; 1 pixel = 0.49 μm) and thresholded at an intensity of 7.5 × 10^-4^ to generate a binary tumor mask. A pixel was included in the tumor mask if it exceeded the PanKRT intensity threshold, was occupied by a segmented cancer cell, or both. This two-criterion definition ensures that genuine tumor regions with locally low PanKRT signals due to staining variability are retained in the tumor compartment mask if annotated cancer cells are present. Connected tumor regions smaller than 7,000 pixels (∼1,670 μm^2^; ∼0.66% of an image) were removed and holes up to 1,000 pixels (∼238 μm^2^) were filled to produce continuous nest regions.

Unique cancer nests were identified by applying connected component analysis to the binary cancer compartment masks. In this framework, two cancer cells are considered members of the same nest if their segmentation footprints overlap with the same connected region of the pixels in the tumor compartment. As a result of the two-criterion definition of a tumor nest, cancer cells separated by small intercellular gaps within a continuous compartment were grouped into the same cancer nest without requiring direct cell-to-cell membrane contact. Each connected component was assigned a globally unique integer identifier numbered sequentially across all the samples. Segmented cancer cells were assigned to the cancer nest where they shared the greatest pixel overlap. In cases where a cancer cell’s segmentation footprint spanned more than one connected nest region, the cell was assigned to the nest occupying the majority of its pixel area. No minimum nest size threshold was applied.

### Cancer nest clustering

Cancer nests were clustered into three groups based on their subtype composition using k-means clustering (k = 3) on the nest-level proportions of classical, intermediate, and basal cancer cells. The resulting clusters were designated classical-rich, intermediate-rich, and basal-rich based on their dominant cancer cell subtypes.

The number of cancer nests per field of view per patient is tabulated (**Supplementary Fig. S2**). The number of nest clusters (k) was selected by maximizing the average silhouette width at k = 2–8. To confirm that the nest-level ECAD association was not driven by patients contributing many nests, we fit a linear mixed-effects model (lme4/lmerTest) of scaled area-normalized ECAD with nest type as a fixed effect and the patient as a random intercept. The significance of nest type was assessed by a likelihood-ratio test against the intercept-only model, and pairwise nest-type contrasts were estimated with emmeans using Bonferroni adjustment **Supplementary Fig. S2D**).

### Distance-based spatial feature extraction

For each unique cancer nest, we defined concentric distance bands extending outward from the nest border: 0–25 μm, 25–50 μm, 50–100 μm, 100–200 μm, 200–300 μm, 300–400 μm, and greater than 400 μm. Distance calibration was performed using a pixel-to-micrometer conversion factor of 500/1024 μm/pixel. Stromal cells were assigned to distance bands based on the Euclidean distance from each cell centroid to the nearest cancer nest border. Within each distance band, the cell type densities, cell type ratios, functional marker positivity rates, Shannon diversity index for immune cell populations, and tumor-stromal ratios were computed. A total of 46 extracellular matrix features were extracted from the COL1A1 fiber segmentation masks within each distance band.

### Proximal versus distal enrichment analysis

For each nest type, we compared spatial features between the proximal (0–50 μm) and distal (greater than 200 μm) zones. Log2 fold-changes were computed as log2(proximal/distal) for each feature. Statistical significance was assessed using t-tests with Benjamini-Hochberg FDR correction. Concordance analysis compared proximal enrichment profiles between basal-rich and classical-rich nests by plotting log2 fold-changes against each other, with features classified as basal-rich-selective, classical-rich-selective, concordantly enriched, or concordantly depleted based on quadrant position.

### QUICHE spatial niche analysis

To identify enriched cellular neighborhoods proximal to cancer nests using an independent spatial method, we applied Quantitative Identification of Cell Heterogeneity and Enrichment (QUICHE) (21). QUICHE defines local cellular niches based on the composition of each cell’s k-nearest neighbors, constructs a niche similarity graph, groups niches into neighborhood categories, and tests for differential spatial enrichment of each neighborhood type between proximal and distal zones relative to cancer nests. The spatial FDR was computed to account for multiple spatial comparisons. Separate QUICHE analyses were performed for basal-rich, classical-rich, and intermediate-rich cancer nests.

### ECM feature extraction and analysis

The ECM organization was characterized using automated collagen fiber segmentation. COL1A1 channel images were processed through TWOMBLI (The Workflow Of Matrix BioLogy Informatics) to generate fiber segmentation masks (22). From each segmentation mask, 46 ECM features were extracted, including the fiber orientation statistics, curvature statistics, fiber length percentiles, organizational metrics, and network properties. ECM features were computed globally per FOV and within local sliding windows for spatial analysis.

### Bulk RNA-seq cohorts

Two independent bulk RNA-seq cohorts were used for the transcriptomic ecotype analysis. The discovery cohort consisted of 277 PDAC patients (**Supplementary Table S3**) profiled by whole-transcriptome RNA sequencing (16). The validation cohort consisted of 147 PDAC patients from The Cancer Genome Atlas (TCGA) Pancreatic Adenocarcinoma (PAAD) project with available RNA-seq and survival data (17).

### EcoTyper analysis

EcoTyper was applied to the OHSU discovery cohort (N = 277) to discover co-occurring cell state patterns (ecotypes) (16). To ensure that established basal and classical lineage markers were included in cell state discovery, we modified the EcoTyper workflow to identify cell states directly from cell-type-specific gene subsets of the bulk expression matrix for each of the 12 cell types, we extracted curated marker gene sets from the pan-cancer EcoTyper reference and subset the bulk expression matrix to the corresponding genes (26,27). The reference gene set was augmented with 19 additional basal and classical lineage markers (GATA6, KRT18, HNF4A, TFF1, REG4, ANX10, CLDN18, LGALS4, DDC, SLC40A1, CLRN3, KRT5, KRT17, S100A2, GPR87, KRT6A, BCAR3, PTGES, and C16orf74). Genes with zero expression across all the samples were removed. Each cell-type-specific expression subset was then subjected to EcoTyper’s non-negative matrix factorization step to decompose samples into transcriptionally defined cell states. To filter the discovered cell states, we recovered each state in two independent PDAC scRNA-seq datasets, the HTAN PDAC cohort (16) and the Werba et al. cohort (23), and retained only states with a recovery Z-score ≥ 1.65 in at least one of the two datasets. The remaining high-confidence cell states were used for cross-cell-type co-occurrence analysis, which identified 11 ecotypes (E1–E11) in the discovery cohort. Ecotype assignments were independently recovered in the TCGA PAAD validation cohort (N = 147) using the cell state signatures and ecotype definitions established in the discovery cohort.

### Survival analysis

Overall survival was assessed using Kaplan–Meier analysis with log-rank tests for pre-specified group comparisons. For each ecotype, hazard ratios were estimated using Cox proportional hazards regression comparing patients assigned to each ecotype versus all others, with 95% confidence intervals reported for all estimates. Prognostic ecotypes were identified by directional concordance of their hazard ratios across the two independent cohorts (discovery and TCGA validation), a replication-based criterion; concordantly protective (E1, E11) and concordantly harmful (E7, E9) ecotypes were carried forward for transcriptomic characterization. The primary survival comparison between the resulting good- and poor-prognosis ecotype groups was assessed by Kaplan–Meier analysis with a log-rank test. Per-ecotype hazard ratios with unadjusted and Benjamini–Hochberg–adjusted p-values are provided in **Supplementary Table S4**.

### MHC class II and gene signature analysis

The expression of MHC class II genes (HLA-DRA, HLA-DPA1, HLA-DPB1, CD74) and co-stimulatory molecules (CD80 and CD86) was compared between the good-prognosis (E1 and E11) and poor-prognosis (E7 and E9) ecotype groups using Wilcoxon rank-sum tests. To account for differences in macrophage abundance, expression values were normalized to the EcoTyper-estimated macrophage fraction (summed abundance of the Monocyte/Macrophage cell states). Basal and classical gene signature scores were computed for each EcoTyper-defined epithelial cell state using published gene sets.

### Statistical analysis

All statistical tests were two-sided. Multiple comparison corrections were applied using Bonferroni correction for pairwise cell-level comparisons and Benjamini-Hochberg FDR correction for feature-level analyses. Bonferroni correction was applied to the small number of pairwise nest-type comparisons to strictly control family-wise error for our primary hypothesis tests, while Benjamini-Hochberg FDR correction was applied to the larger feature-level spatial screens (∼250 features) to balance sensitivity and specificity. Statistical significance was set at p < 0.05. All the analyses were performed using R (version 4.x) and Python (version 3.x).

## Data and code availability

All code for image analysis, spatial feature extraction, and statistical analysis is available at https://github.com/cameronwalk14/PDAC_MIBI. Processed MIBI-TOF imaging data, cell segmentation files, and derived single-cell and spatial feature tables were deposited on Zenodo under DOI https://doi.org/10.5281/zenodo.21108550. Bulk RNA-seq data from TCGA PAAD cohort are publicly available through the NCI Genomic Data Commons.

## Authors’ Contributions

Conception and design: C. Walker, M. Angelo, T. Risom, A. Gentles, R. Sears, and C. Pelz. Acquisition of data: H. Piyadasa. Analysis and interpretation of data: C. Walker, R. Wang, B. Benard, J. Eng, and K. Hawthorne. Writing, review, and revision: All authors. Study supervision: M. Angelo, T. Risom, R. Sears, and A. Gentles.

## Acknowledgements

We would like to thank all the patients. We thank P. Gupta, S Qi, and S. Hassan for their thoughtful discussions related to this work. This work was supported by National Institutes of Health grants U54 CA209971 (A.G), U01 CA264611 (A.G), U54CA20997105 (M.A.), 5DP5OD01982205 (M.A.), 1R01CA24063801A1 (M.A.), 5R01AG06827902 (M.A.), 5UH3CA24663303 (M.A.), 5R01CA22952904 (M.A.), 1U24CA22430901 (M.A.), 5R01AG05791504 (M.A.), 5R01AG05628705 (M.A.), the Department of Defense W81XWH2110143 (M.A.), the Wellcome Trust and other funding from the Bill and Melinda Gates Foundation, Cancer Research Institute, the Parker Center for Cancer Immunotherapy and the Breast Cancer Research Foundation. The funders had no role in the study design, data collection, and analysis, decision to publish or manuscript preparation. The authors used a generative AI assistant (Claude, Anthropic) to support language editing in the manuscript. All AI-assisted text was reviewed, edited, and verified by the authors, who take full responsibility for the accuracy and integrity of the final content.

---

**Supplementary Figure S1.**
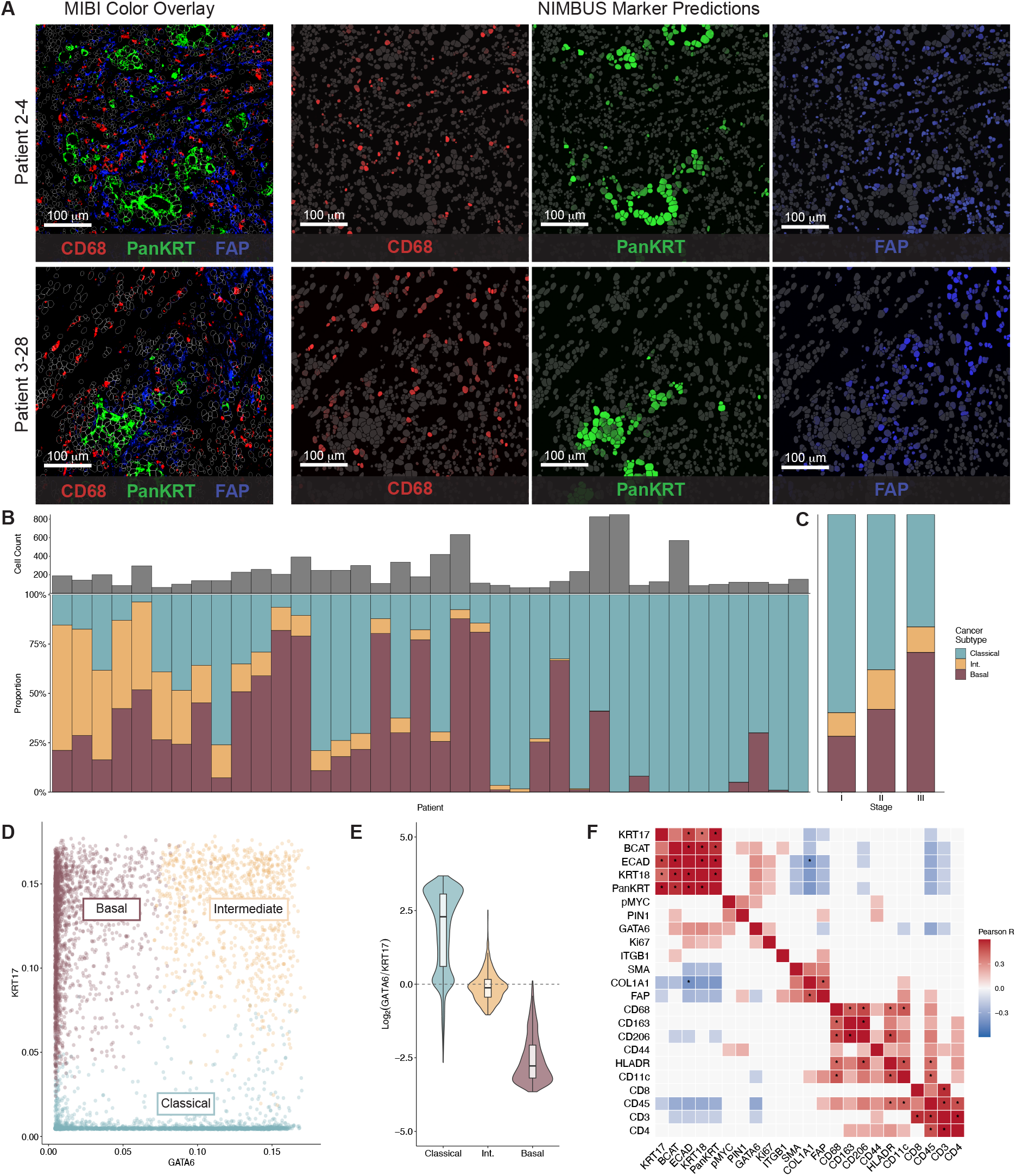
Cell type annotation validation and cancer subtype distribution. (A) Representative MIBI-TOF color overlay with NIMBUS prediction maps for CD68, Pan-Keratin, and FAP. (B) Cancer cell subtype composition by patient across the MIBI cohort. (C) Cancer cell subtype composition by stage. (D) Single-cell co-expression of GATA6 and KRT17. (E) Log2-transformed GATA6-to-KRT17 marker expression by cancer subtype. (F) Pearson correlation heatmap of marker expression across all cells.

**Supplementary Figure S2.**
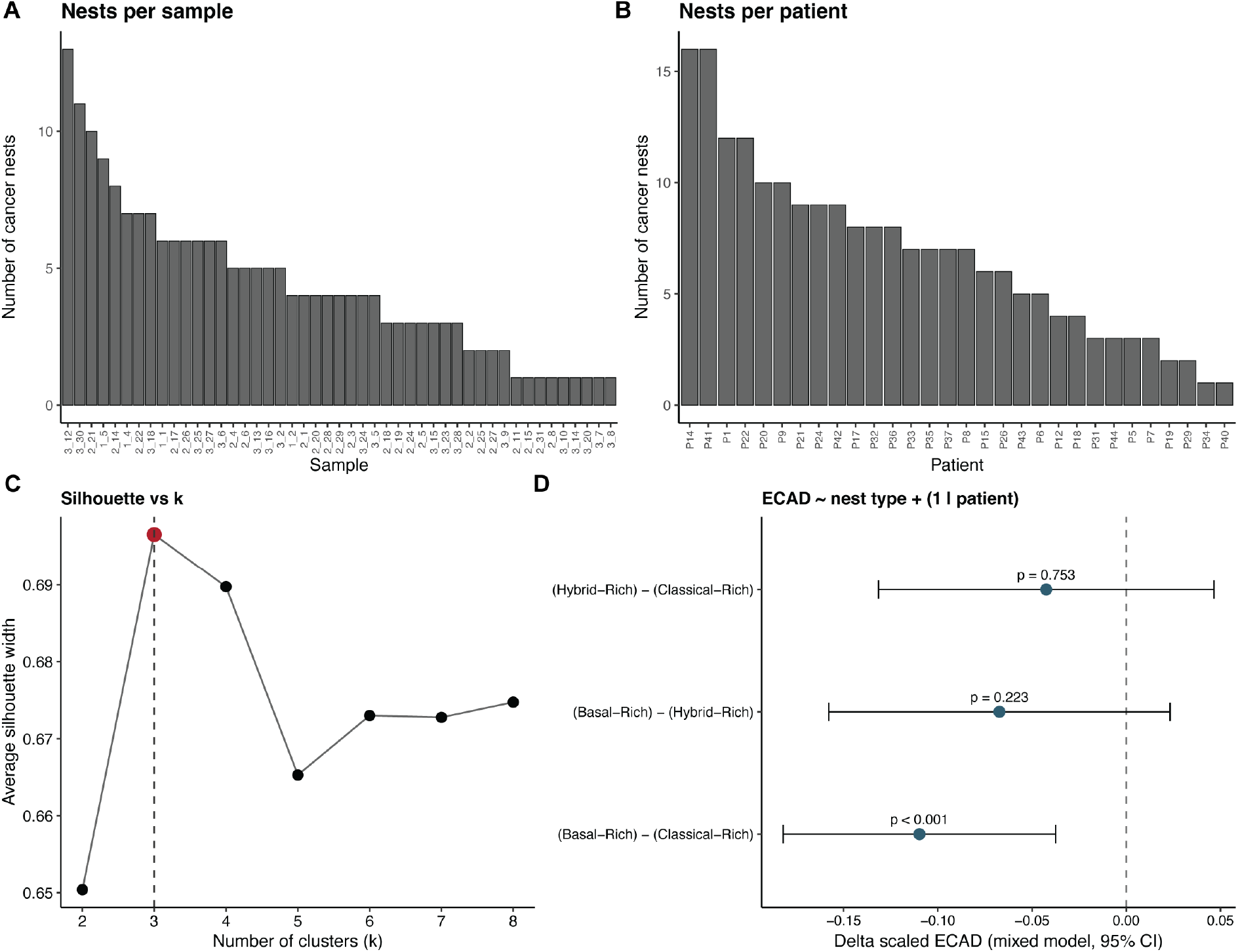
Cancer-nest sampling depth, cluster-number selection, and patient-level modeling of epithelial ECAD. (A) Number of spatially contiguous cancer nests per MIBI-TOF field of view; each bar is one sample (B) Number of cancer nests per patient; each bar is one patient with at least one cancer nest. (C): Average silhouette width for k-means clustering of nests by cancer-subtype composition across candidate cluster numbers k = 2–8. The average silhouette width is maximized at k = 3 (highlighted point; dashed line). (D) Linear mixed-effects model of area-normalized E-cadherin (ECAD) expression as a function of nest type with patient as a random intercept (ECAD ∼ nest type + (1 | patient)). Points and horizontal bars denote the estimated pairwise differences in scaled ECAD between nest types with 95% confidence intervals; the dashed line marks no difference. Between-patient variation accounted for 38% of the total variance (intraclass correlation = 0.38). Nest type remained a significant predictor of ECAD after accounting for patient (likelihood-ratio test χ²₂ = 13.5, P = 1.2 × 10⁻³), and Classical-Rich nests retained significantly higher ECAD than Basal-Rich nests (Bonferroni-adjusted P = 9.6 × 10⁻⁴).

**Supplementary Figure S3.**
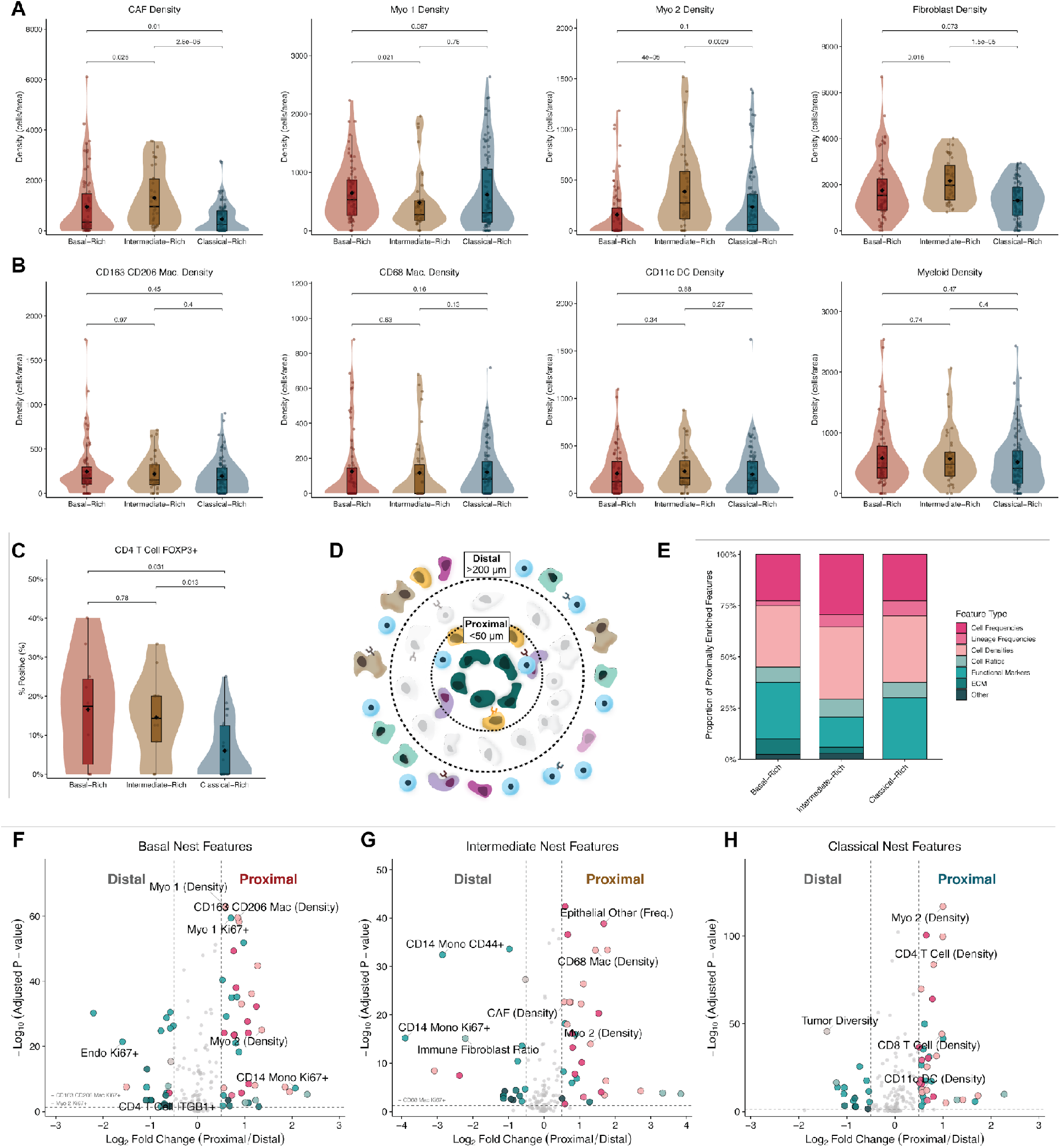
Fibroblast and myeloid cell densities, regulatory T cell frequency, and proximal enrichment by cancer nest type. (A) Fibroblast densities proximal (0–50 µm) to each nest type: CAF density, Myo 1 density, Myo 2 density, and total fibroblast density. (B) Myeloid cell densities proximal to each nest type: CD163/CD206 macrophage density, CD68 macrophage density, CD11c dendritic cell density, and total myeloid density. (C) Percentage of FOXP3+ CD4 T cells (Tregs) proximal to each nest type. (D) Visualization of proximal vs. distal enrichment analysis. (E) Proportion of feature type enriched near each nest type. (F-H) Volcano plots showing proximal versus distal enrichment of spatial features for Basal-Rich (F), Intermediate-Rich (G), and Classical-Rich (H) nests, highlighting enrichment of cancer-associated fibroblast subsets near Basal-Rich nests. Violin plots show per-nest values with pairwise Wilcoxon rank-sum tests; boxplots indicate median and interquartile range with mean shown as diamond.

**Supplementary Figure S4.**
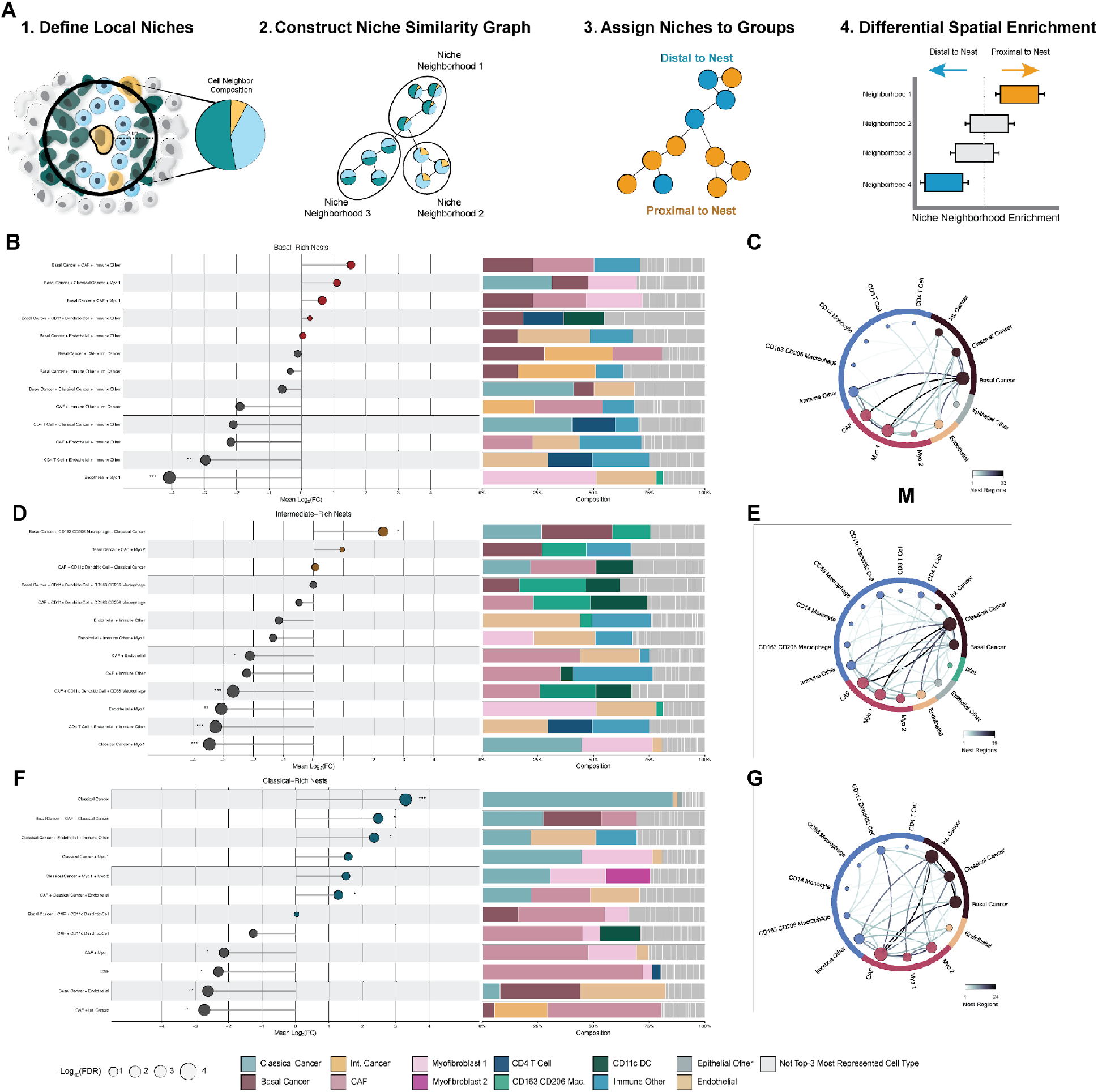
QUICHE spatial niche enrichment analysis. Cleveland dot plots showing mean log2 fold-change for significantly enriched or depleted neighborhoods (left) with corresponding niche type composition (right) and cell type co-occurrence network diagrams for Basal-Rich, Intermediate-Rich, and Classical-Rich cancer nests.

**Supplementary Figure S5.**
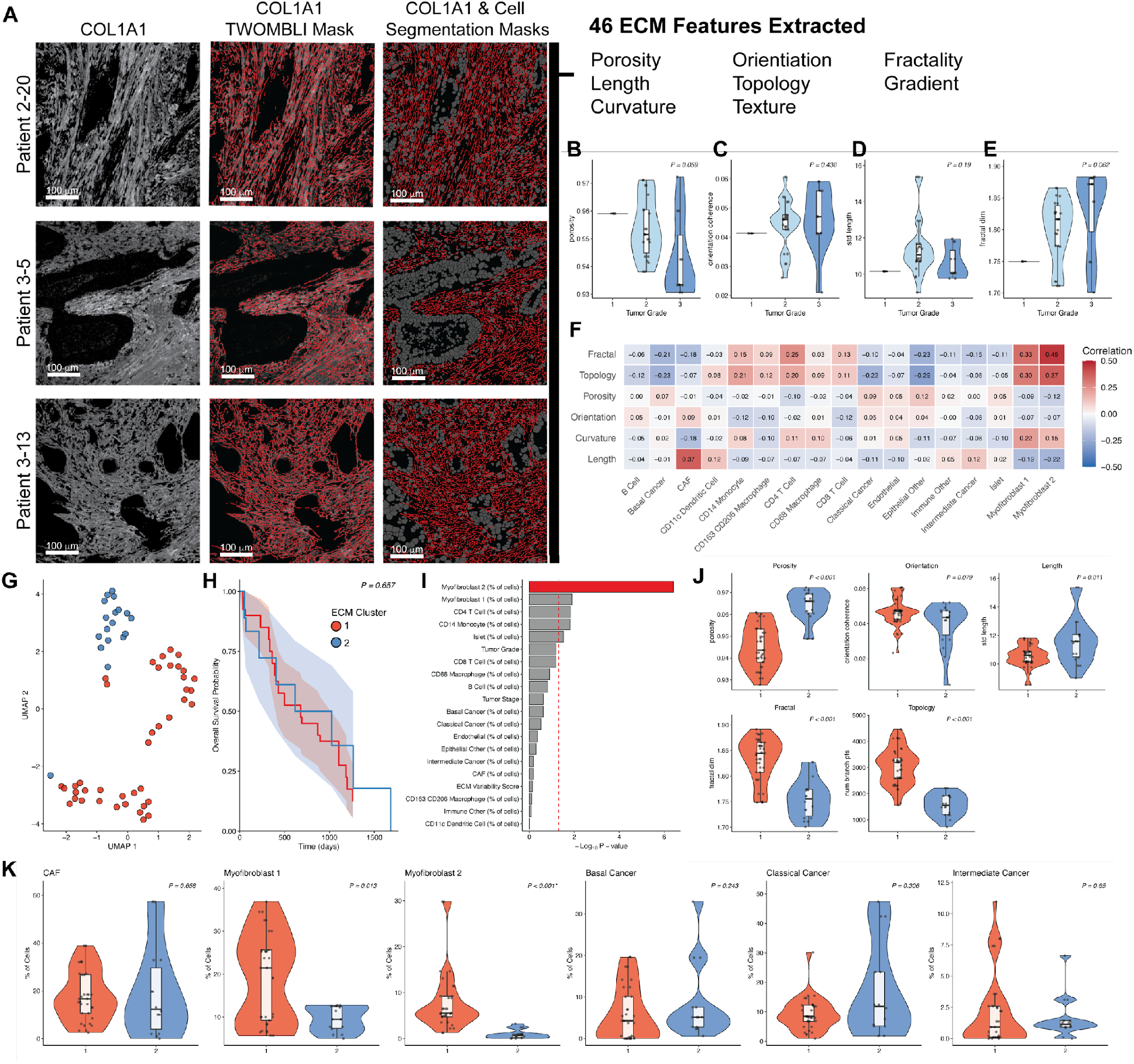
Collagen fiber segmentation and ECM feature analysis. (A) Representative COL1A1 images, TWOMBLI-generated segmentation masks, and overlay with cell masks. (B–E) ECM features by tumor grade: porosity (B), orientation coherence (C), fiber length variability (D), fractal dimensionality (E). (F) Spearman correlations between ECM feature categories and cell type frequencies. (G) UMAP projection of ECM profiles by cluster. (H) Kaplan-Meier survival by ECM cluster. (I) Association strength of clinical and biological variables with ECM cluster membership; red indicates statistically significant associations. (J) ECM features by cluster. (K) Cell type frequencies by ECM cluster.

**Supplementary Figure S6.**
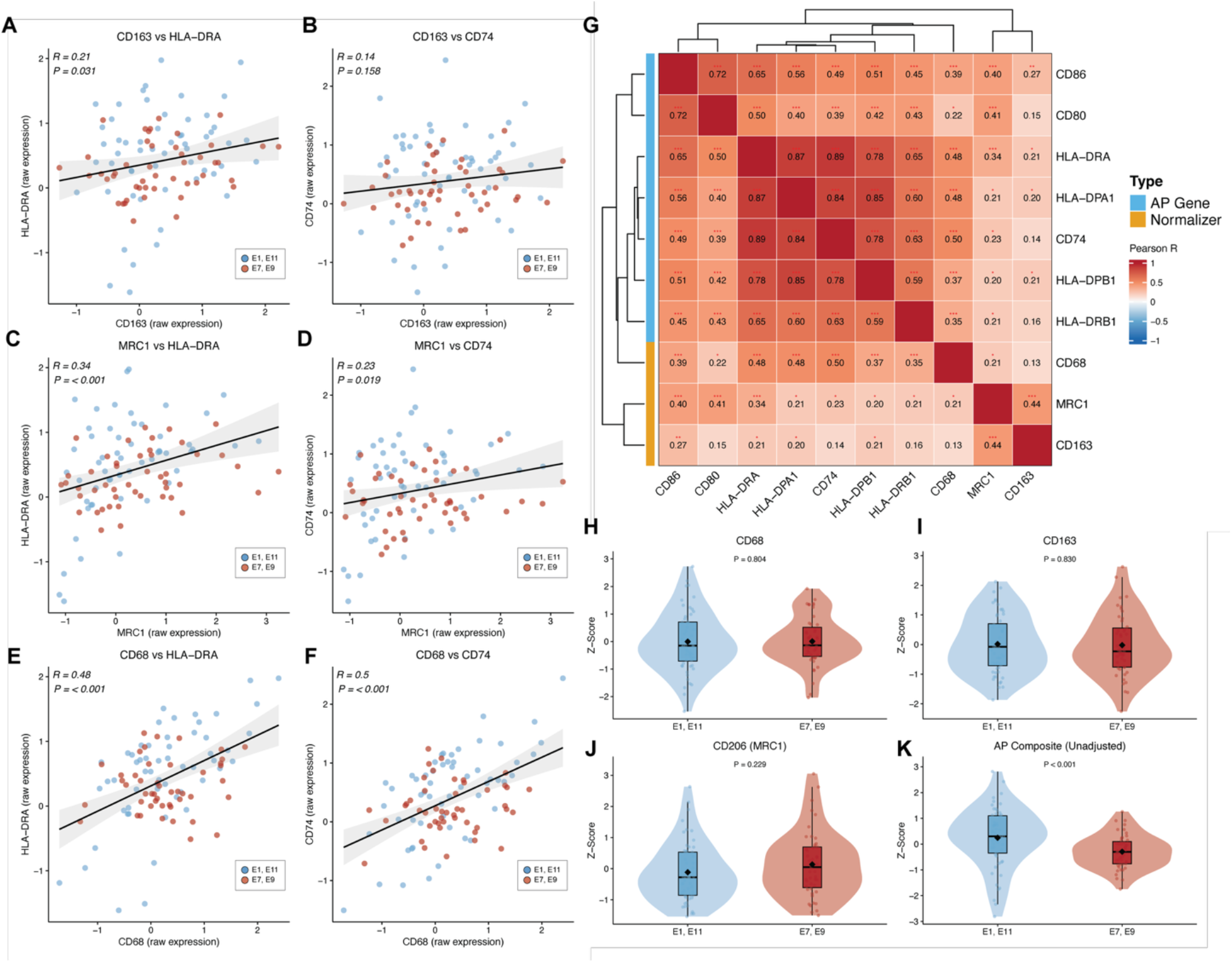
Antigen presentation differences between prognostic ecotypes are not explained by macrophage abundance. (A–F) Pearson correlation scatter plots between myeloid marker genes and antigen presentation genes in the EcoTyper-deconvoluted myeloid compartment. (G) Pearson correlation heatmap of myeloid normalizer genes (CD68, CD163, MRC1) and antigen presentation (AP) genes across all samples. Hierarchical clustering separates AP genes from CD163 and MRC1, while CD68 clusters adjacent to the AP gene block. Significance indicated by asterisks (*P < 0.05, **P < 0.01, ***P<0.001). (H-K) Expression of myeloid marker genes and composite antigen presentation score between concordant good-prognosis (E1, E11) and poor-prognosis (E7, E9) ecotypes.

**Supplementary Figure S7.**
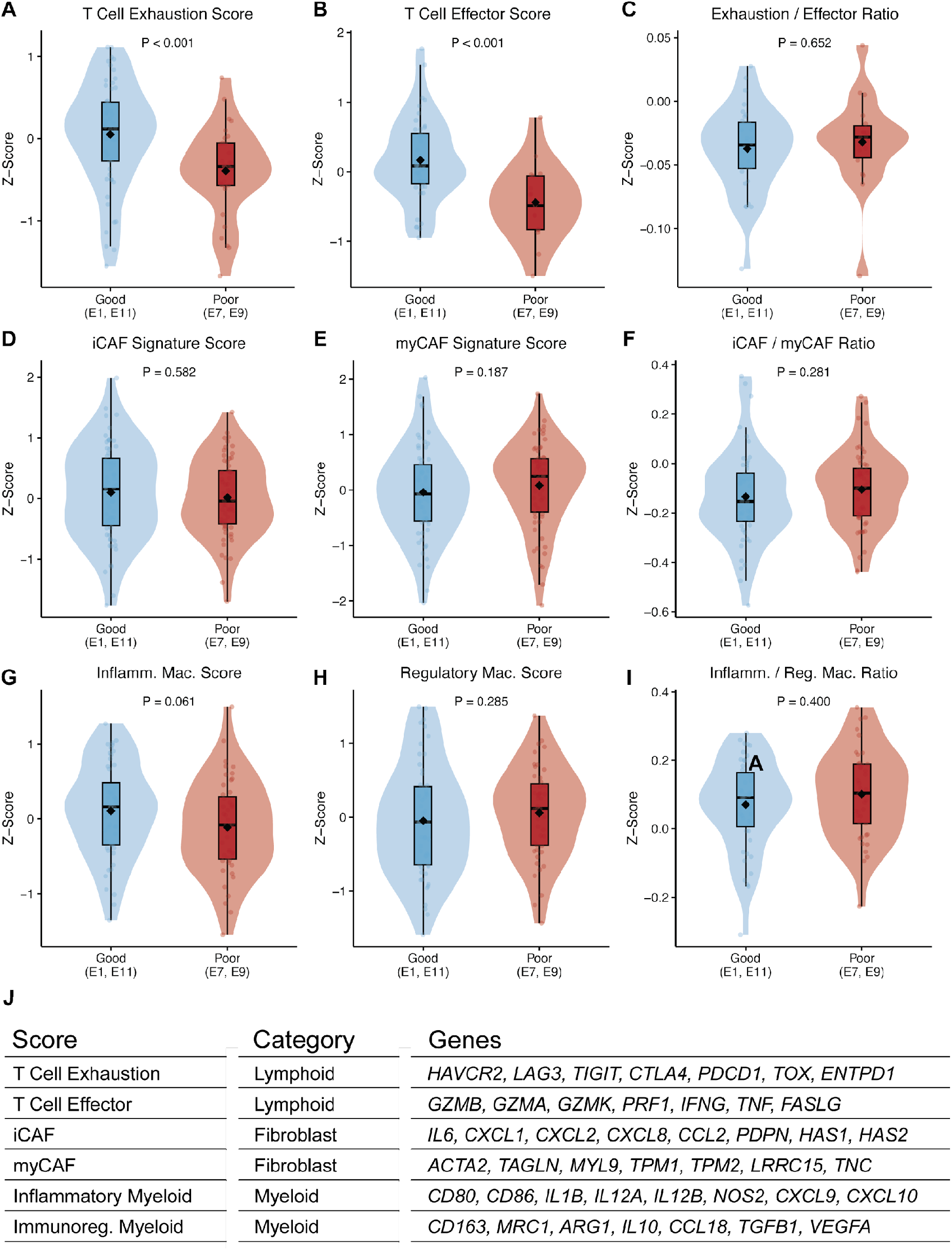
Gene signature scores across prognostic EcoTyper ecotypes. (A–C) T cell exhaustion (A), effector (B), and exhaustion/effector ratio (C) comparing good versus poor prognosis ecotypes. (D–F) iCAF (D), myCAF (E), and iCAF/myCAF ratio (F). (G–I) Inflammatory macrophage (G), regulatory macrophage (H), and inflammatory/regulatory ratio (I). (J) Gene signatures used for scoring.

**Supplementary Table S1.** Clinicopathologic characteristics of the MIBI-TOF cohort (n = 34 patients). Values are n (%) of patients; unknown values shown separately.

| Characteristic | Category | n (%) (N=34) |
| --- | --- | --- |
| **Sex** | Female | 20 (59%) |
|  | Male | 14 (41%) |
| **Age at diagnosis** | ≤70 years | 23 (68%) |
|  | >70 years | 11 (32%) |
| **AJCC stage** | I | 6 (18%) |
|  | II | 20 (59%) |
|  | III | 3 (9%) |
|  | Unknown/NA | 5 (15%) |
| **Grade** | 1 | 1 (3%) |
|  | 2 | 13 (38%) |
|  | 3 | 6 (18%) |
|  | Unknown/NA | 14 (41%) |
| **Neoadjuvant treatment** | No | 20 (59%) |
|  | Yes | 10 (29%) |
|  | Unknown/NA | 4 (12%) |
| **PurIST subtype** | Classical | 25 (74%) |
|  | Basal-like | 9 (26%) |
| **Vital status at last follow-up** | Alive | 7 (21%) |
|  | Deceased | 27 (79%) |

**Supplementary Table S2.** MIBI-TOF antibody panel. Metal-conjugated antibodies used for MIBI-TOF imaging, listed by mass channel.

| Mass channel | Target | Clone |
| --- | --- | --- |
| 69Ga | Calprotectin | MAC387 |
| 71Ga | Chymase/Tryptase | EPR13136/EPR9522 |
| 113In | VIM | D2H13 |
| 115In | SMA | SP171 |
| 141Pr | PIN1 | G-8 |
| 142Nd | TCF1/7 | C63D9 |
| 143Nd | CD4 | EPR6855 |
| 144Nd | CD14 | D7A2T |
| 145Nd | FAP | ERP20021 |
| 146Nd | FoxP3 | 236A/E7 (abcam) |
| 147Sm | Phospho-c-MYC | 4B12 |
| 148Nd | CD31 | EP2095 |
| 149Sm | Collagen Hybridizing Peptide-biotin | CHP-biotin |
| 150Nd | ECAD | 24E10 |
| 151Eu | CD56 | MRQ42 |
| 152Sm | CD209 | DCN46 |
| 153Eu | CD206 | 685645 |
| 154Sm | GATA6 | D61E4 |
| 155Gd | CD74 | D5N3I |
| 156Gd | CD68 | D4B9C |
| 157Gd | CD44 | E7K2Y |
| 158Gd | CD8 | C8144B |
| 159Tb | CD3 | D7A6E |
| 160Gd | Amylase | D55H10 |
| 161Dy | CD11c | EP1347Y |
| 162Dy | ITGB1 | D2E5 |
| 163Dy | CD163 | D6U1J |
| 164Er | Beta Catenin | D10A8 |
| 165Ho | Fibronectin | E5H6X |
| 166Er | KRT18 | E431-1 |
| 167Er | CD20 | L26 |
| 168Er | Synaptophysin | D8F6H |
| 169Tm | CD45 | D9Mal |
| 170Er | Podoplanin | D2-40 |
| 171Yb | COL1A1 | E8F4L |
| 172Yb | CD9 | D3H4P |
| 173Yb | PanCK | AE1/AE3 |
| 174Yb | KRT17 | SP95 |
| 175Lu | Ki67 | 8D5 |
| 176Yb | HLADR | EPR3692 |

**Supplementary Table S3.** Clinicopathologic characteristics of the OHSU discovery RNA-seq cohort (289 tumor samples from 277 patients). Values are n (%) of patients; The discovery cohort comprises 289 whole-transcriptome RNA-seq samples from 277 patients. PurIST subtype from bulk RNA classification; grade recorded for a subset.

| Characteristic | Category | n (%) (N=289) |
| --- | --- | --- |
| Sample type | Primary | 218 (75%) |
|  | Metastatic | 71 (25%) |
| Specimen site | Pancreas | 218 (75%) |
|  | Liver | 29 (10%) |
|  | Peritoneum | 18 (6%) |
|  | Lung | 7 (2%) |
|  | Other | 17 (6%) |
| PurlST molecular subtype | Classical | 225 (78%) |
|  | Basal-like | 64 (22%) |
| Histologic grade | Well | 19 (7%) |
|  | Moderate | 109 (38%) |
|  | Poor | 37 (13%) |
|  | Not recorded | 124 (43%) |

**Supplementary Table S4.** Per-ecotype survival hazard ratios across the discovery and validation cohorts. Cox proportional hazards ratios (HR) with 95% confidence intervals comparing overall survival between patients assigned versus not assigned to each ecotype (assignment analysis) or in the top 25% versus bottom 25% of ecotype abundance (abundance analysis) in the OHSU discovery and TCGA-PAAD validation cohorts. P, unadjusted log-rank/Wald p-value; P(BH), Benjamini–Hochberg–adjusted p-value within each cohort × analysis family (11 ecotypes). Prognostic ecotypes were selected by directional concordance of HRs across cohorts rather than by individual significance.

**Assignment analysis (assigned vs not assigned to ecotype)**
| Ecotype | Discovery HR (95% CI) | Discovery P | Discovery P(BH) | Validation HR (95% CI) | Validation P | Validation P(BH) |
| --- | --- | --- | --- | --- | --- | --- |
| E1 | 0.88 (0.62–1.25) | 0.467 | 0.642 | 0.64 (0.38–1.10) | 0.108 | 0.325 |
| E2 | 0.90 (0.59–1.36) | 0.619 | 0.744 | 1.57 (0.63–3.91) | 0.329 | 0.741 |
| E3 | 1.69 (1.13–2.53) | 0.011 | 0.099 | 0.83 (0.26–2.65) | 0.755 | 0.917 |
| E4 | 1.00 (0.57–1.75) | 0.993 | 0.993 | 0.96 (0.41–2.21) | 0.917 | 0.917 |
| E5 | 1.82 (1.09–3.03) | 0.022 | 0.099 | — | — | — |
| E6 | 0.91 (0.58–1.42) | 0.676 | 0.744 | 1.08 (0.57–2.05) | 0.807 | 0.917 |
| E7 | 1.36 (0.97–1.90) | 0.070 | 0.155 | 1.05 (0.58–1.92) | 0.866 | 0.917 |
| E8 | 0.39 (0.14–1.05) | 0.061 | 0.155 | — | — | — |
| E9 | 1.52 (0.87–2.67) | 0.143 | 0.263 | 1.09 (0.44–2.71) | 0.848 | 0.917 |
| E10 | 0.78 (0.52–1.18) | 0.242 | 0.380 | 2.28 (1.31–3.98) | 0.004 | 0.033 |
| E11 | 0.53 (0.30–0.93) | 0.027 | 0.099 | 0.16 (0.02–1.21) | 0.076 | 0.325 |

**Abundance analysis (top 25% vs bottom 25% of ecotype abundance)**
| Ecotype | Discovery HR (95% CI) | Discovery P | Discovery P(BH) | Validation HR (95% CI) | Validation P | Validation P(BH) |
| --- | --- | --- | --- | --- | --- | --- |
| E1 | 1.11 (0.78–1.59) | 0.549 | 0.750 | 1.41 (0.75–2.65) | 0.291 | 0.741 |
| E2 | 0.86 (0.60–1.24) | 0.424 | 0.750 | 0.77 (0.41–1.46) | 0.428 | 0.741 |
| E3 | 0.91 (0.64–1.31) | 0.630 | 0.750 | 0.90 (0.48–1.67) | 0.742 | 0.742 |
| E4 | 1.02 (0.71–1.45) | 0.928 | 0.928 | 0.76 (0.41–1.42) | 0.389 | 0.741 |
| E5 | 1.08 (0.76–1.53) | 0.681 | 0.750 | 1.18 (0.63–2.20) | 0.606 | 0.741 |
| E6 | 1.09 (0.76–1.56) | 0.638 | 0.750 | 0.70 (0.38–1.29) | 0.252 | 0.741 |
| E7 | 0.91 (0.64–1.29) | 0.584 | 0.750 | 0.90 (0.50–1.62) | 0.721 | 0.742 |
| E8 | 1.25 (0.88–1.77) | 0.215 | 0.750 | 0.83 (0.46–1.50) | 0.531 | 0.741 |
| E9 | 1.50 (1.04–2.15) | 0.029 | 0.319 | 0.84 (0.46–1.53) | 0.573 | 0.741 |
| E10 | 0.92 (0.64–1.32) | 0.648 | 0.750 | 0.62 (0.34–1.12) | 0.113 | 0.741 |
| E11 | 0.87 (0.61–1.24) | 0.448 | 0.750 | 1.22 (0.68–2.20) | 0.502 | 0.741 |

